# Highly Parallelized, Multicolor Optogenetic Recordings of Cellular Activity for Therapeutic Discovery Applications in Ion Channels and Disease-associated Excitable Cells

**DOI:** 10.1101/2022.04.11.487888

**Authors:** Gabriel B Borja, Hongkang Zhang, Benjamin N Harwood, Jane Jacques, Jennifer Grooms, Romina Chantre, Dawei Zhang, Adam Barnett, Christopher A Werley, Yang Lu, Steven Nagle, Owen B McManus, Graham T Dempsey

**Affiliations:** Q-State Biosciences, Cambridge, MA 02139; USA

**Keywords:** optogenetics, Optopatch, high throughput screening, ion channels, voltage-gated sodium channels, pain, human iPSC models, drug discovery

## Abstract

Optogenetic assays provide a flexible, scalable and information rich approach to probe compound effects for ion channel drug targets in both heterologous expression systems and associated disease relevant cell types. Despite the potential utility and growing adoption of optogenetics, there remains a critical need for compatible platform technologies with the speed, sensitivity, and throughput to enable their application to broader drug screening applications. To address this challenge, we developed the Swarm™, a custom designed optical instrument for highly parallelized, multicolor measurements in excitable cells, simultaneously recording changes in voltage and calcium activities at high temporal resolution under optical stimulation. The compact design featuring high power LEDs, large numerical aperture optics, and fast photodiode detection enables all-optical individual well readout of 24-wells simultaneously from multi-well plates while maintaining sufficient temporal resolution to probe millisecond response dynamics. The Swarm delivers variable intensity blue-light optogenetic stimulation to enable membrane depolarization and red or lime-light excitation to enable fluorescence detection of the resulting changes in membrane potential or calcium levels, respectively. The Swarm can screen ~10,000 wells / day in 384-well format, probing complex pharmacological interactions via a wide array of stimulation protocols. To evaluate the Swarm screening system, we optimized a series of heterologous optogenetic spiking HEK293 cell assays for several voltage-gated sodium channel subtypes including Nav1.2, Nav1.5, and Nav1.7. The Swarm was able to record pseudo-action potentials stably across all 24 objectives and provided pharmacological characterization of diverse sodium channel blockers. We performed a Nav1.7 screen of 200,000 small molecules in a 384-well plate format with all 560 plates reaching a Z’≥0.5. As a demonstration of the versatility of the Swarm, we also developed an assay measuring cardiac action potential and calcium waveform properties simultaneously under paced conditions using human induced pluripotent stem (iPS) cell-derived cardiomyocytes as an additional counter screen for cardiac toxicity. In summary, the Swarm is a novel high-throughput all-optical system capable of collecting information-dense data from optogenetic assays in both heterologous and iPS cell-derived models, which can be leveraged to drive diverse therapeutic discovery programs for nervous system disorders and other disease areas involving excitable cells.

## 1 Introduction

The development of effective therapeutics for disorders of the nervous system has been challenging in comparison with other areas such as cancer and cardiovascular disease (Schoepp, 2011). Therapeutics for neurological disorders historically have had longer development periods and lower FDA approval rates compared to most other indication areas (Kaitlin, 2014; Dowden and Munro, 2019). For example, chronic pain is a neurological condition affecting more than one hundred million people in the United States and remains a major unmet medical need (Institute of Medicine of the National Academies, 2011). Individuals suffering from chronic pain often face significant loss of productivity and a decrease in overall quality of life (Connor, 2009). Opioid treatments, which are the most frequently prescribed treatment for chronic pain, have considerable undesirable side effects (Benyamin et al., 2008). The addictive properties of opioids create a substantial risk for the development of abuse disorders that can further negatively impact quality of life for patients. Existing non-opioid alternative drugs for chronic pain are less efficacious for a large subset of patients (Grosser et al., 2017). Despite the massive unmet medical and societal need, the pharmaceutical industry has yielded few new therapeutics for chronic pain patients and has similarly struggled for other nervous system disorders. As such, there remains a critical need for new technologies and approaches to support development of novel therapeutics for chronic pain and other neurological diseases with improved efficacy and safety profiles (Gribkoff and Kaczmarek, 2017).

Ion channels represent a promising class of drug targets for a variety of nervous system-based disorders (Waszkielewicz et al., 2013), as they are essential components in signal generation and propagation in neurons and other electrically excitable cell types. The characterization of pathogenic mutations in human ion channels has directly linked ion channel function to human disease pathologies, including ‘channelopathies’ associated with severe genetic epilepsies, chronic pain conditions, and many others (Snowball and Schorge, 2015; Imbrici et al., 2016; Albury et al., 2017; Reid et al., 2017). Channel biophysical properties across the hundreds of ion channel targets have been deeply investigated, though the mechanisms that connect genetic and pharmacological modulation of ion channel properties to systems level outcomes in patients remains an area of active research. Bridging this knowledge gap is critical in the identification of effective therapeutics that target different ion channels. Despite the continued advancements in understanding ion channel properties, it remains a challenge to translate these findings to effective therapeutics.

One class of ion channels with strong genetic evidence for involvement in disease pathologies and broad recognition as important drug targets in the pharmaceutical industry are the voltage-gated sodium (Nav) channels (Wang et al., 2017). Nav channels are transmembrane proteins that are expressed and localized to the membranes of electrically excitable cells, such as neurons, and are responsible for the selective transport of Na^+^ ions. There are nine members of the Nav channel class (Nav1.1-1.9), each with critical roles in diverse tissue types (de Lera Ruiz and Kraus, 2015). For example, Nav1.1, 1.2, 1.3, and 1.6 are expressed in the central nervous system, and their dysfunction in these channels can lead to epilepsy, migraine, autism, and ataxias (Ogiwara et al., 2009). Nav1.7, 1.8, and 1.9 are expressed predominantly in dorsal root ganglion (DRG) sensory neurons, and dysfunction leads to pain disorders such as small fiber neuropathy (Eijkenboom et al., 2019) and congenital insensitivity to pain (Cox et al., 2006). Nav1.5 is expressed in heart muscle, and pathogenic mutations affecting this channel lead to Brugada syndrome, long QT syndrome, and atrial fibrillation (Keating and Sanguinetti, 2001; Splawski et al., 2002; Veerman et al., 2015). Our understanding of Nav channel biology has advanced dramatically over the last decade, however drug development for these targets remains a major challenge. Achieving the ideal molecular profile that balances selectivity, state dependence and the preferred pharmacological activity in intact disease relevant cell types like neurons or cardiomyocytes is a key factor that has ultimately prevented the full realization of effective therapeutics for this class of targets.

A major challenge within Nav channels and, more generally, in ion channel drug discovery efforts is the limitations of the existing technologies, assays, and disease models, which lack either the throughput or the information content required to derive necessary insights into complex human biology and pharmacological responses that enable translation to efficacious therapies. Consequently, there remains a need to develop model systems that are both information-rich and high-throughput to enable rapid identification and optimization of novel drug templates. While methods such as manual patch clamp electrophysiology (E-phys) provide rich information content, high resolution, and flexibility to fine-tune assay conditions, this approach is severely limited by throughput in the context of drug discovery screening applications. Automated E-Phys development has made significant strides in achieving higher throughput than manual patch clamp, but still has lower throughput and is less affordable than optical assays, which are key considerations when conducting primary screening of large chemical libraries. First generation optical based technologies such as FLIPR and E-VIPR (Schroeder, 1996; Huang et al., 2006) are cost effective and well adapted for high throughput screens (HTS), however they do not maintain high temporal resolution of single millisecond scale required to faithfully record electrical spiking, thus limiting information provided on complex drug mechanisms. In addition, most plate-based fluorescence assays rely on non-physiological stimuli that can alter pharmacological sensitivity and impact translation to therapeutic efficacy in humans. Given the limitations of current technologies, there is a clear need for new neuroscience drug discovery platforms applicable to ion channel targets that are capable of screening hundreds of thousands to millions of compounds while maintaining physiologically relevant assay conditions and providing information-rich data comparable to manual or automated patch clamp methods.

A promising solution to addressing current challenges in ion channel drug discovery lies in recent advances in optogenetic methodologies (Song and Knöpfel, 2016; Dempsey and Werley, 2017; Zhang and Cohen, 2017), which can enable high resolution all-optical electrophysiology achieving both dramatic improvements in throughput and deep information content measurements of excitable cells. One approach, called Optopatch™ (Hochbaum et al., 2014), relies on genetically engineered optogenetic actuators such as the channelrhodopsin variant CheRiff that can be expressed in a variety of cell types to initiate cell depolarization and subsequent action potential (AP) initiation using a blue light stimulus (488nm). Changes in membrane potential can be quantified with exquisite sensitivity using either a voltage sensitive fluorescent protein such as QuasAr (637nm) (Hochbaum et al., 2014) or a voltage sensitive dye such as BeRST1 (635nm) (Huang et al., 2015) by recording the changes in red fluorescence over time. When a blue-light driven actuator and red-light fluorescence voltage reporter are co-expressed, all-optical electrophysiology can be conducted with high spatial and temporal resolution. The achievable spatial and temporal resolution of optogenetic electrophysiology now enable the development of high throughput imaging systems to support drug discovery (Werley et al., 2017).

To translate the advances in optogenetic actuator and sensor technology to a robust, all-optical therapeutic discovery platform that can be leveraged for diverse drug targets such as ion channels we have developed the Swarm™, a novel high-throughput screening instrument for optogenetic assays. Here, we describe the physical and optical characteristics of the Swarm instrument. The Swarm can deliver blue-light stimuli across entire individual wells in a multi-well plate, eliciting changes in membrane potential and then recording cell depolarizations and firing via red-light excitation and fluorescence detection. To enable the high-throughput capacity of Swarm, the instrument stimulates and records from 24 wells simultaneously, which allows information from an entire 96-well or 384-well plate to be collected in 4 or 16 recordings, respectively. The system allows for optogenetic pacing and analysis of electrically excitable cell types in a multi-wavelength format with four input excitation channels (three independent wavelengths and one for patterned illumination) and three independent fluorescence channels for multicolor recording. We have also developed custom software to control the instrument and a suite of tools for automated data analysis.

To demonstrate the utility of the Swarm for ion channel drug discovery, we applied a set of custom optogenetic assays using HEK293 cell lines that heterologously express an optogenetic actuator and a series of voltage gated-sodium channel (Nav1.x) subtypes including Nav1.2, Nav1.5, and Nav1.7 (herein referred to as ‘Nav1.x spiking HEK assays’) and are compatible with a fluorescent voltage sensing dye BeRST1 (Zhang et al., 2016, 2020). In humans, Nav1.7 is a critical modulator of pain sensitivity and is expressed primarily in sensory neurons, playing a central role in neuronal action potential initiation(Gingras et al., 2014; Hameed, 2019). Nav1.7 is a genetically validated target for novel pain therapeutics with known loss- and gain-of-function mutations identified in humans that lead to dramatic hypo- or hyper-sensitivity to pain, respectively (Goldberg et al., 2007; Emery et al., 2016; Shields et al., 2018). Given the genetic validation of Nav1.7 as a drug target, significant effort has been devoted to identifying small molecule inhibitors that block Nav1.7. However, to date, none have demonstrated sufficient clinical efficacy to gain approval for pain related indications. Several challenges have hindered the development of small molecule inhibitors including a requirement for subtype selectivity to avoid adverse effects in the cardiovascular and nervous system resulting from block of other sodium channel subtypes such as Nav1.5 and Nav1.2, respectively (Mulcahy et al., 2019). Nonselective voltage-gated sodium blockers, such as lidocaine have clinical utility and are frequently used as local anesthetics, however systemic administration can cause side effects such as dizziness, shortness of breath, and nausea (Cherobin and Tavares, 2020). Compounds with high Nav1.7 subtype selectivity, such as PF-05089771, have shown limited efficacy in clinical pain studies, possibly due to a strong state-dependent blocking mechanism, lack of blood brain barrier penetrance, and poor performance in sensory neurons (McDonnell et al., 2018). These issues highlight a need to identify new classes of Nav1.7 inhibitors that meet the required characteristics to enable their use for treating chronic pain.

Using Nav1.x spiking HEK assays (Zhang et al., 2016, 2020) in a 384-well plate format, we show validation of the Swarm for high-throughput optogenetic screening. First, we demonstrate assay performance through uniform signal detection across all 24 detection channels and across whole 384-well control plates formatted with alternating columns of positive and negative control compounds. Next, we use an extensive panel of control pharmacology across Nav1.2, Nav1.5, and Nav1.7 spiking HEK assays to show that measured IC_50_ values and state dependence metrics are consistent with literature reports. We then use the Swarm to perform a HTS screen of 200,000 small molecules to identify novel inhibitors of Nav1.7. The demonstrated Z’ (>0.5) and on-plate quality control metrics (IC_50_ of tetracaine) demonstrate high performance of the instrument and assay. We further report on potency, state dependence, and selectivity of confirmed hits from the screen along with tool compounds which are either clinically used or that have failed in clinical trials as comparators. Lastly, we show that the Swarm can also be used to record action potentials and calcium transients simultaneously under paced conditions in human iPS cell-derived cardiomyocytes along with control pharmacological validation, enabling utilization of the Swarm instrument for counter screening for Nav channel programs or more broadly for screening applications in human cellular models.

## 2 Materials and Methods

### 2.1 Swarm Instrument Design

The Swarm read head objective modules are made of custom-designed, CNC-machined, anodized aluminum (Protolabs, Maple Plain, MN). The 12-objective read heads are built as six independent 2-objective modules that hold the lenses, dichroics, and filters. The compact design of the read head module is dictated by the pitch between wells of a 96-well plate (9mm) and complies with the optical design to maximize the light throughput, minimize stray light, and maximize positioning accuracy. A spring-loaded plate pusher is designed to hold 96-, 384-, or 1536-well microplates. The plate pusher arm is articulated above the read-heads mounted flush to the top deck to provide smooth translation through a low-profile X-Y linear stage (Thorlabs, Newton, NJ), so that all wells can be measured, 24 wells at a time.

The Swarm read head comprises two banks of 12 objective modules, 24 in total, mounted to the top deck beneath the well plate. Each objective module houses more than 40 compact optics (lenses, filters, masks), LEDs, and photodiodes. Within the illumination path, there are 4 independent channels with high power LEDs (Luxeon, Alberta, Canada) – a red channel (627nm), a lime channel (567nm), an unpatterned blue channel (470nm), and a patterned blue channel (470nm with a chrome mask at the conjugate image plane). Light is delivered to the sample using a series of high numerical aperture aspheric relay lenses (LightPath, Orlando, FL) in 4-f configuration. The lens relay configuration was simulated and optimized with optical design software (Zemax, Kirkland, WA) to ensure maximum light delivery efficiency and to inform read head mechanical manufacturing tolerance. The designed focus spot size is 1mm^2^ at the sample after the final objective lens. A red (637 nm), lime (563 nm), or blue (470 nm) bandpass filter (Semrock, Rochester, NY) is placed in front of each LED to narrow to the targeted excitation wavelengths. In the detection path, three output color channels are separated by dichroic mirrors (Semrock, Rochester, NY) detected simultaneously by three independent photodiodes (Hamamatsu, Hamamatsu, Japan): one photodiode is behind a red filter (736 nm) for a voltage indicator, one behind an orange filter (600 nm) for a calcium indicator, and one photodiode is behind a GFP filter (520 nm) for protein expression detection or other measurements.

We designed and fabricated custom-made printed circuit boards to hold the essential electronics components, collect/transmit signals, and deliver power to the Swarm. An LED board and 2 photodiode boards are attached to each two-objective module. Each LED board contains 8 high-power LEDs (2 x 4 LEDs per objective). The two photodiode boards contain 6 photodiodes (2 x 3 photodiodes per objective) and six independent picoammeter circuits that amplify the photodiode signal. Each two-objective module is also connected to a signal/driver board. The signal/driver board has two functions. First, it contains the LED driver circuitry that can deliver up to 1 A of low-noise constant current to the high-power LEDs in the LED board. Second, it collects the amplified signals from the photodiode boards and delivers them to a Compact Data Acquisition (DAQ) system (National Instruments, Austin, TX), which provides synchronized analog input for the 72 total photodiode channels and synchronized analog output control for the 96 LEDs. Each signal/driver board is connected to a power distribution bus board that delivers stable DC voltage from high-performance rugged power supplies (Acopian, Easton, PA).

### 2.2 Nav1.x Spiking HEK Cell Culture

Spiking HEK cells were used for Swarm instrument validation, the 200K small molecule screen, and tool compound pharmacological characterization. The spiking HEK cell lines consisted of HEK293 cells stably expressing Nav.1.7, Nav1.5, or Nav1.2 co-expressed with the blue-shifted channelrhodopsin actuator CheRiff-EGFP. Spiking HEK cell lines and culture conditions are as previously described with slight modifications (Zhang et al., 2016, 2020). In brief, vials of Nav1.x spiking HEK cells were thawed and plated at 3 million cells/10cm dish. Cell culture growth media contained DMEM, 10% FBS, 1% GluMax (Gibco, Waltham, MA) and MEM Non-Essential Amino Acids (ThermoFisher, Waltham, MA). To expand total cell numbers required for HTS, cells were grown at 37 °C with 5% CO_2_ for 48 hours to approximately 70% confluence. After 48 hours, 4mL of cell culture medium was removed and 300μL of Kir2.1 lentivirus was added to the 10cm dish. Cells were incubated for 24 hours to facilitate Kir2.1 lentivirus transduction. The following day, 384-well plates (789836, Greiner Bio-One) were treated with Poly-D-lysine (20μg/mL) for 1 hour at 37°C and aspirated. Kir2.1 transduced HEK293 cells were then trypsin treated (0.5%), resuspended, centrifuged, washed, and resuspended in fresh DMEM with serum. Cells were then replated in Poly-D-lysine treated 384-well plates at a density of 30,000 cells per well. Cells were then incubated at 37 °C overnight to achieve 100% confluence prior to running spiking HEK assays.

### 2.3 Nav1.x Spiking HEK Assays

Imaging was performed on Nav1.x spiking HEK cells 24 hours after plating on 384-well plates. Prior to compound addition, medium was removed from cells using a flick and tap method, and then 1 μM BeRST dye was added using a Janus Mini liquid handling robot (PerkinElmer, Waltham, MA). The cells were then incubated for 30 minutes at 30°C with no exposure to light. Following incubation, the cells were washed using Tyrodes solution (containing, 125mM NaCl, 8mM KCl, 3mM CaCl_2_, 1mM MgCl_2_, 30mM glucose, 10mM HEPES, pH 7.35) and compounds were added using a Janus Mini liquid handling robot (PerkinElmer, Waltham, MA). Compounds were prepared in 1 μL spots on a separate compound plate at a concentration of either 666-fold or 1666-fold over screening concentration, with negative controls (DMSO) and positive controls (1 μM TTX), as wells as a reference compound (Tetracaine in a 3-fold dilution series) on each plate. Following compound addition, plates were incubated for an additional 40 minutes at room temperature in the dark.

Each plate was read in two orientations throughout the studies to ensure data redundancy, first with well A1 designated as the top left corner of the plate, and then immediately again with well P24 as the top left corner. Instantaneous quality control metrics were generated using custom MATLAB (MathWorks, Natick, MA) software following each two-orientation plate read. Figures included heatmaps, signal-to-noise ratio, a list of traces indicating potentially problematic wells, and tetracaine concentration response curves (CRCs) with calculated IC_50_ values to ensure that the plate did not need to be imaged again due to user or instrument error. For Nav1.2 and Nav1.5 spiking HEK assays, imaging was performed using a process nearly identical to Nav1.7, with minor differences in potassium bath concentration and stimulus frequency (see Results section for details).

### 2.4 Compound Libraries

Several small molecule libraries were used for instrument validation and our HTS campaign. For instrument validation, the commercially available Prestwick Chemical library™ (Prestwick Chemical Inc., Illkirch-Graffenstaden, France) was used. This library consists of 1280 1520 small molecules, most of which are FDA approved, prepared in 384-well plates at a stock concentration of 10mM in DMSO. The 200,000 small molecule library screened for Nav1.7 inhibitors was compiled from a series of commercially available sources and internally assembled with three goals: diversity, CNS drug-like properties, and chemical scaffolds amenable to medicinal chemistry. Compound handling was performed using Janus and Janus Mini automated liquid handling instruments (PerkinElmer, Waltham, MA).

### 2.5 Nav1.x Spiking HEK Assay Data Analysis

For each well, the time course of fluorescence intensity was collected from a single objective and the collected raw trace was corrected for photobleaching by dividing the raw intensity time trace by the median filtered intensity. To reduce well-to-well variability, the photobleaching corrected trace was normalized to the steady state height of fluorescence plateau reached during the 500 ms long blue pulse (TP9), which is mainly determined by CheRiff depolarization, stable, and insensitive to sodium channel inhibitors. Next, for Nav1.7 and Nav.1.5, the peak amplitude during each blue light test pulse (TP) was extracted and the mean spike amplitude across all test pulses was used to calculate overall compound potency. For Nav1.2, mean spike areas under TP1, TP8, and TP10 were used to calculate compound potency. Since different regions within the same 384-well plate were imaged by different objectives, to further reduce the variability across different objectives, at the beginning of each imaging day, a sentinel plate with DMSO and a positive control compound (TTX for Nav1.2 and Nav1.7; tetracaine for Nav1.5) was imaged to establish the baseline and magnitude of Nav channel dependent signal from each objective. For all the following screening plates imaged on the same day, the extracted parameters from each well were normalized to the mean values of positive control and negative control wells imaged by the same objective from the sentinel plate. Since each plate was imaged using the regular orientation and the second orientation by rotating the plates by 180 degrees, each well was imaged by a pair of objectives and the mean values from these two objectives was used for next step of analysis. After objective-based normalization, to reduce plate-to-plate variability, each calculated parameter was further normalized based on the positive control wells and negative control wells from the same plate. For each plate, to evaluate assay quality, a Z’ factor was calculated as Z’ = 1−3(σ_p_+σ_n_)/|(μ_p_-μ_n_)|, where σ_p_ is the standard deviation of all positive control wells, σ_n_ is the standard deviation of all negative control wells, μ_p_ is the mean of the positive controls and μ_n_ is the mean of the negative controls.

### 2.6 Cardiomyocyte Tissue Culture, Imaging and Data Analysis

Human iPS cell-derived cardiomyocytes (Fujifilm Cellular Dynamics, Madison, WI) were cultured in clear 96-well Greiner COC plates which were plasma treated and coated with 0.1% gelatin (STEMCELL Technologies, Vancouver, Canada). iCell cardiomyocytes plating medium from the iCell Cardiomyocytes Media Kit was used to plate the cardiomyocytes at 60,000 cells/well and then exchanged with iCell Cardiomyocytes maintenance medium after 24 hours. Four days after plating, the hiPSC-derived cardiomyocytes were transduced with lentiviral expression vectors for the blue light actuator CheRiff and the calcium indicator jRGECO1a. After 24 hours, the medium was removed and exchanged for iCell maintenance medium to remove the virus. Medium (150 μL) was exchanged with fresh maintenance medium every 48 hours. The cardiomyocytes imaging experiments were performed 9 days after plating.

Prior to imaging, maintenance medium was removed from the cells using a multichannel pipette, and then 150 μL of 1 μM BeRST1 dye was added to each well. The cells were incubated with BeRST1 dye for 30 minutes at 30°C in the dark. Following incubation, BeRST1 dye was washed out using 4 mM K^+^ Tyrode’s solution. The cells were incubated with the Tyrode’s solution and preliminary recordings were made on the Swarm instrument with the enclosure temperature set at 30°C. Compounds were then added in a dilution series to three replicate wells per concentration. The compounds were incubated for 10 minutes before the second post-drug addition recordings were made.

For the cardiomyocyte imaging, two separate photodiodes were used to simultaneously record both the BeRST1 voltage fluorescence and the jRGECO1a Ca^2+^ fluorescence. The photodiode voltage recordings were collected with an NI-9202 Analog Input module (National Instruments, Austin, TX) at 2000 Hz. The raw voltage recordings were low-pass filtered with a moving average filter and then bleach corrected to report the change in fluorescence (dF/F). Blue light stimulus artifacts were removed by linear interpolation between the data points immediately before and after the blue stimulus pulse. Post-drug addition traces were normalized to the maximum dF/F of the predrug addition traces for the same well. The max AP amplitude was extracted from each test pulse epoch and averaged across all 10 blue light test pulses to determine mean spike amplitude. AP70 width was calculated as the time between crossing 30% of maximum fluorescence between the upstroke and downstroke. The AP70 width was only calculated and averaged from epochs where the cardiomyocytes fired APs. The Ca^2+^ transients were quantified by computing the integrated area under the curve of the photobleach corrected jRGECO1a traces and averaged over all 10 test pulse epochs. Error bars represent the standard error of the mean (SEM) across three replicate wells each. Significance testing was performed using a one-way ANOVA with Dunnett’s test and p values were considered significant if <0.05. For the simultaneous voltage and calcium imaging experiments, the significance testing was conducted using a one-tailed t-test and p values were considered significant if <0.05.

## 3 Results

### 3.1 Swarm for Highly-parallelized, Multicolor Optogenetic Recordings

We designed and built the Swarm instrument, which records both voltage and calcium activity in 24 wells simultaneously under optical simulation. The read head contains 24 objective modules, which enable flexibility in validation testing, optogenetic imaging, troubleshooting, and repair. Each objective module has over 40 optical elements comprised of low-cost molded aspheric lenses, low-noise photodiodes, high-power LEDs, and high-quality thin film filters and dichroics. Within each module, there are four excitation LED channels (637nm red, 563nm lime, 470nm blue, and patterned 470nm blue) optically cleaned by excitation filters, and three emission channels (736nm, 573nm, and 520nm) collected by photodiode detectors (Figure 1A). The objective modules are aligned in parallel to form two banks of twelve modules that comprise the whole 24-objective read head. Each assay scan reported for this study requires approximately 9 seconds (3 seconds for imaging, 6 seconds of overhead, e.g. DAQ initialization, stage moving to read new wells, saving recordings). Imaging a 384-well plate requires 16 consecutive scans (~2.5 minutes) using the read pattern shown in Figure 1B. We constructed custom printed circuit boards attached to each objective module as shown in Figure 1C: (1) a LED board containing four high power LEDs for the four excitation optical paths (red, lime, blue, and patterned blue). (2) Two photodiode boards containing three photodiodes and three independent picoammeter circuits to measure and amplify the signal collected in the three emission channels (QuasAr, RFP, and GFP). The overall Swarm instrument and superstructure footprint on an 18” by 24” optical breadboard is shown in Figure 1D.

**Figure 1.**
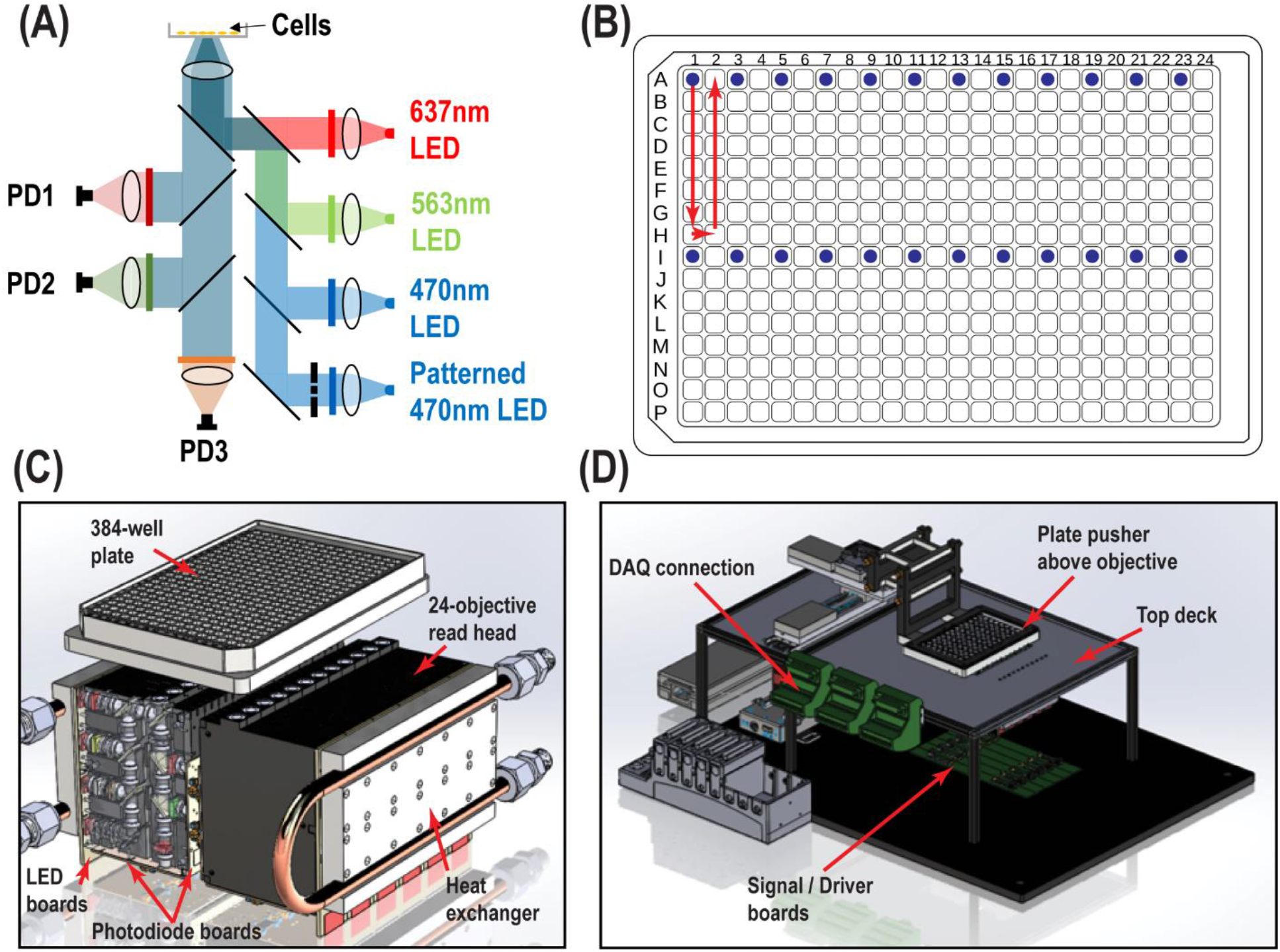
Swarm instrument schematics. (A) Each objective module contains four excitation LED paths (637nm, 563nm, 470nm, and patterned 470nm) and three emission channels (736nm, 573nm, and 520nm), which are collected by photodiode detectors. Chrome masks are placed on conjugate image planes to enable patterned blue light stimulation. Beam shaping optics not shown. (B) 384-well plate and objective modules illuminating twenty-four wells (blue dots) and scan pattern (red arrow) to image the entire plate. (C) Each individual objective module has two banks of twelve modules that comprise the whole 24-objective read head. Each module contains multiple lenses, filters and masks to deliver light from the LED board to the bottom of the well. The signal amplification picoammeters measure currents from the sample which are collected by each photodiode channel. Water cooled heat exchangers are fastened to the LED boards. (D) Surrounding superstructure of Swarm. A plate pusher arm articulates a 384-well plate over a 24-objective read head, which is fastened below the top deck. The DAQ connectors hang to the side of the top deck rails.

### 3.2 Optopatch Nav1.7 Spiking HEK Swarm Assay Validation

Nav1.x spiking HEK cells can fire sodium channel dependent pseudo-APs which afford robust signals reflecting Nav channel function and pharmacology. The basic components of Nav1.x spiking HEK cells are shown in Figure 2A. Nav1.x spiking HEK cells stably expressed a Nav1.x channel, the target of interest, and a modified channelrhodopsin (Zhang et al., 2016, 2020). In addition, an inwardly rectifying potassium channel (Kir2.1) was transiently expressed by lentiviral transduction. Kir channel expression was required to hyperpolarize the resting membrane potential in order to maintain Nav channels in a non-inactivated state and enable repolarization following stimulation. Expression of Kir channels also allowed tunable control of resting potential by adjusting extracellular potassium concentrations in imaging (Dai et al., 2008). For the primary Nav1.7 assays, we used 8 mM bath potassium to achieve a resting membrane potential of approximately −70 mV, which is near the physiological value in native sensory neurons (Liu et al., 2017). Cells were loaded with a bright far-red fluorescent membrane potential dye, BeRST1 prior to imaging to provide a rapid and linear fluorescence readout of membrane voltage, which is compatible with red LED intensity levels from Swarm (4W/cm^2^). With this model, cells can be depolarized upon blue light stimuli leading to Nav1.x channel activation and the firing of pseudo-APs. The AP amplitude can be recorded by measuring BeRST1 fluorescence intensity and is sensitive to Nav channel inhibitor pharmacological modulation. For Nav1.7 spiking HEK cells, a 10 test-pulse protocol was used and the mean spike amplitude across all the test pulses was applied to evaluate Nav1.7 inhibitor potency and efficacy (Zhang et al., 2020). Figure 2B shows all the 16 wells imaged by objective 1 from a zebra sentinel plate with alternating columns of wells containing a Nav1.7 blocker tetrodotoxin (TTX; 1 μM) as a positive control and dimethylsulfoxide (DMSO; 0.15%) as a negative vehicle control. For the single objective, the average waveform amplitudes for the eight DMSO treated wells showed excellent separation from those of the eight TTX treated wells. The 40% signal reduction is due to Nav1.7 inhibition and all residual signal is mediated by voltage actuator CheRiff-mediated depolarization. Since optogenetic assay results can be sensitive to light illumination intensity, we further validated the instrument homogeneity across different objectives by examining the DMSO and TTX traces collected from each objective. Figure 2C shows that after fine tuning of analog output voltages to each blue LED, the Swarm instrument generates a consistent TTX sensitive, Nav1.7 dependent signal amplitude across all the different objectives covering a full 384-well plate. To further reduce the assay variability resulting from subtle differences in LED intensity across different objectives, we also included an objective-wise normalization step in the analysis pipeline (see Methods section). Figure 2D shows a heatmap from a zebra sentinel plate in which DMSO and 1 μM of TTX were placed in alternative columns and no apparent objective or position biased effects were observed.

**Figure 2.**
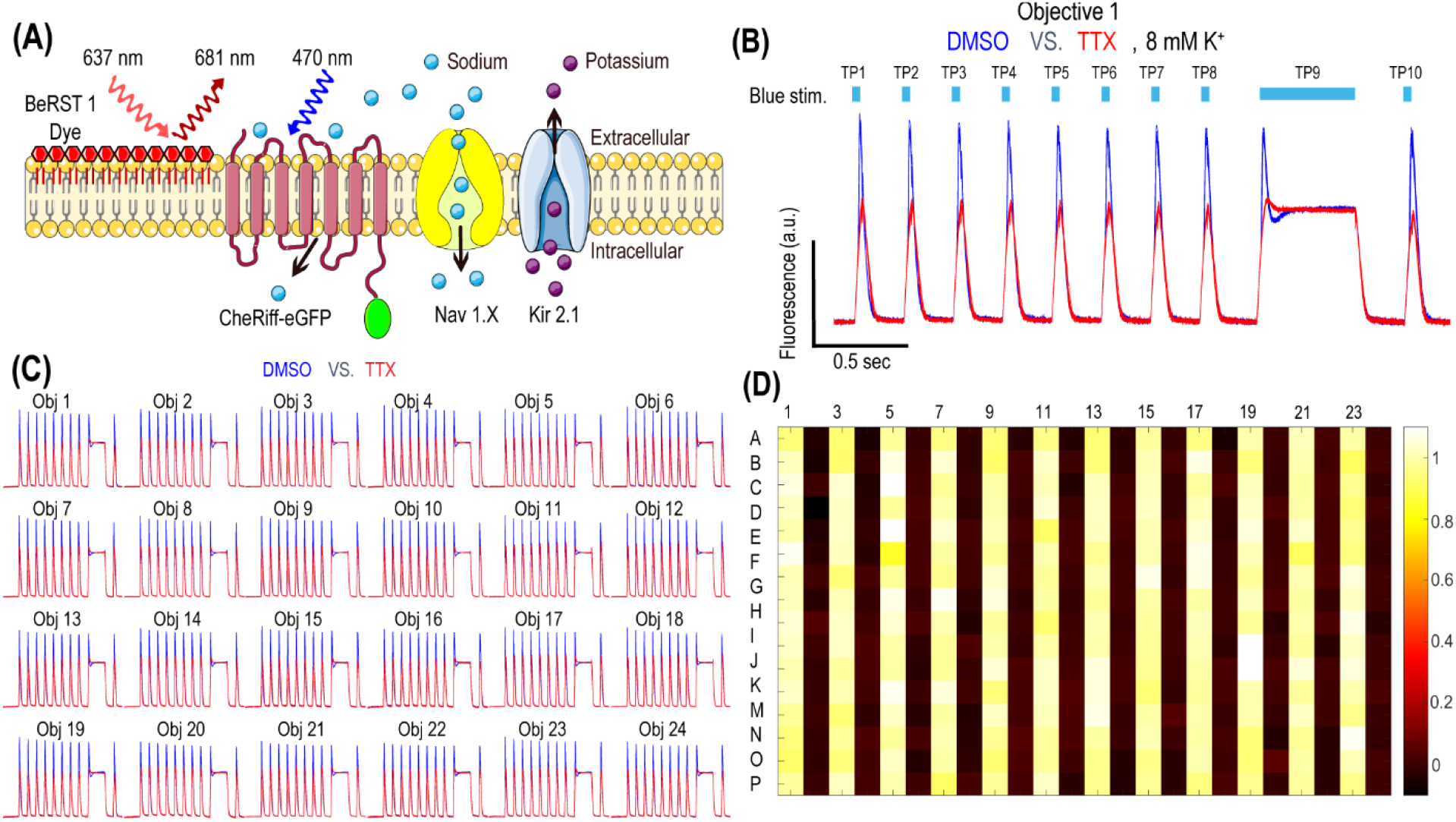
Nav1.7 Spiking HEK assay performance with the Swarm instrument. (A) Diagram of key components in Nav1.x spiking HEK cells, including voltage sensitive dye BeRST1, CheRiff-eGFP, Kir2.1, and the target of interest, Nav1.x channel. (B) Representative fluorescence traces from eight wells of DMSO or 1 μM TTX treated Nav1.7 spiking HEK cells stimulated with a 10 test-pulse blue light stimulation protocol, imaged by Objective 1 of the Swarm instrument. The bath [K^+^] is 8 mM. (C) Fluorescence traces collected from all the 24 objectives. The Nav1.7 spiking HEK cells were treated with either DMSO or 1 μM TTX and were stimulated by the protocol shown in panel (B). Each trace was averaged based on 8 adjacent wells imaged by the same objective. (D) The heat map from a representative Nav1.7 spiking HEK 384-well sentinel plate. All the odd columns were treated with DMSO vehicle control, and all the even columns were treated with 1 μM TTX.

The Optopatch Nav1.7 spiking HEK assay, when run using a serial epifluorescence microscope, was reported to yield IC_50_ values comparable to an automated patch clamp platform such as IonWorks Barracuda (Zhang et al., 2020). The parallel recording feature of Swarm greatly reduces imaging time per plate and subsequently improves overall throughput while maintaining overall assay performance. When a diverse set of compounds were tested using both Swarm and a custom single-well compatible epifluorescence microscope, the instruments yielded IC_50_ values for each compound differing less than two-fold across the two platforms (Supplementary Figure 1).

### 3.3 Tool Compound Pharmacology for Swarm Assay Validation Across Different Nav Channels

In addition to Nav1.7, the Optopatch spiking HEK assay can be applied to study the pharmacology of other Nav subtypes, including Nav1.2 (a major Nav subtype in the brain) and Nav1.5 (dominant Nav subtype in the heart), both of which are important counter screen targets to confirm Nav1.7 selectivity. We established stable cell line expression of Nav1.2/CheRiff and Nav1.5/CheRiff in HEK293 cells and Kir2.1 was introduced to the cells via lentiviral transduction prior to imaging. Similar to the Nav1.7 assay, upon blue light stimulation, Nav1.2 and Nav1.5 spiking HEK cells fire pseudo-APs and the spike amplitude and integrated spike area are sensitive to compound modulation (Figure 3A). The spike width is wider in Nav1.2 and Nav1.5 spiking HEK cells than Nav1.7 spiking HEK cells, possibly due to different Nav subtype channel gating properties or different background potassium current levels in different HEK cell lines (Ponce et al., 2018). Different bath potassium concentrations were used for Nav1.2 (8 mM) and Nav1.5 (6 mM), based on the consideration that the cardiac Nav1.5 subtype has a more hyperpolarized half-inactivation voltage V1/2 than the neuronal subtypes Nav1.2 and Nav1.7 (Vilin et al., 2012; Wang et al., 2015). Adjusting the bath potassium concentration to generate similar levels of channel inactivation across sodium channel subtypes enabled more consistent comparison of pharmacological effects. Like the Nav1.7 assay, we extracted parameters from multiple test pulses to comprehensively evaluate compound effects on Nav1.2 and Nav1.5 channels and then derive overall compound IC_50_ values. We validated the assay sensitivity and accuracy by performing concentration response experiments using a set of 15 Nav tool compounds covering a wide range of subtype specificities, mechanisms of action, and binding sites (Table 1, Figure 3B, Supplementary Figure 2). Under current assay conditions, the nonselective Nav inhibitors amitriptyline, tetracaine, and vixotrigine can inhibit all the three Nav subtypes with similar potency (Figure 3B). The Nav subtype selective compounds JNJ63955918, PF-05089771, and tetrodotoxin demonstrated the expected subtype selectivity consistent with literature reports (Alexandrou et al., 2016; Flinspach et al., 2017; Tsukamoto et al., 2017). Only JNJ63955918 showed more than 50-fold Nav1.7 selectivity against both Nav1.2 and Nav1.5. While both tetrodotoxin and PF-05089771 potently blocked Nav1.7, they lacked or only had modest subtype specificity against Nav1.2 (Figure 3B). In addition to assessing subtype selectivity profiles, Nav1.x spiking HEK assays on Swarm can be used to determine compound mechanism of action. The compound state dependence (Kr/Ki) was defined as the ratio Nav1.7 IC_50_ value at TP1 under 4 mM potassium (Figure 3A) over the IC_50_ value at TP10 under 8 mM potassium (Figure 2B). As expected, compounds that act on the voltage sensor domain IV (e.g. PF-05089771), as local anesthetics (e.g. lidocaine, tetracaine), and as prototypical anticonvulsants (e.g. carbamazepine, lamotrigine) show stronger state-dependent block than pore blockers (e.g. TTX) and peptide blockers (e.g. JNJ63955918).

**Figure 3.**
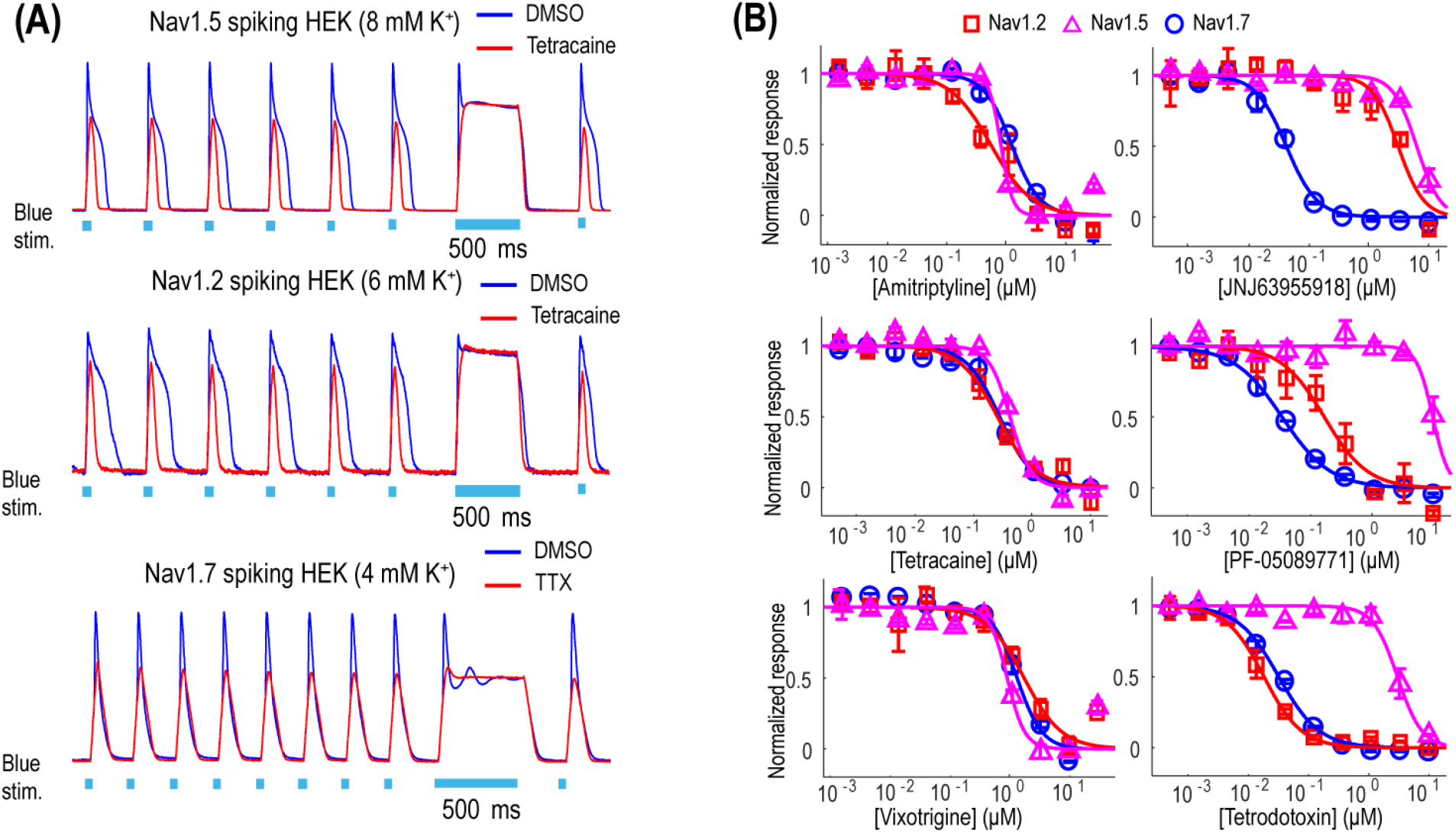
Tool compound concentration-response curves on different Nav channel subtypes. (A) Representative fluorescence traces from DMSO, 1 μM TTX, or 10 μM tetracaine treated Nav1.2 spiking HEK cells (8 mM bath [K^+^]), Nav1.5 spiking HEK cells (6 mM bath [K^+^]), and Nav1.7 spiking HEK cells (4 mM bath [K^+^]). For Nav1.2 and Nav1.5 spiking HEK assays, the cells were stimulated at 2 Hz. Eight test pulses were applied before the 500 ms long pulse and only action potentials triggered by the third to the eighth pulses were plotted for these two Nav channel subtypes; for Nav1.7 spiking HEK cells, the cells were stimulated at 4 Hz and all ten triggered action potentials are plotted. (B) Concentration-response curves of 3 non-selective Nav channel blockers (Amitriptyline, Tetracaine, Vixotrigine) and 3 subtype-selective Nav channel blockers (JNJ63955918, PF-05089771, Tetrodotoxin) on Nav1.2 (8 mM bath [K^+^]), Nav1.5 (6 mM bath [K^+^]), and Nav1.7 (8 mM bath [K^+^]). Error bars indicate SEM based on measurements from 2 wells.

**Table 1:** Tool compound results. Nav1.x IC50 values and state dependence from 15 tool compounds with different potencies and working mechanisms. *State dependence is defined as the ratio of IC_50_ values of Nav1.7 spiking HEK assay at TP1 at 4 mM K^+^ over TP10 IC_50_ values at 8 mM K^+^ (Figure 2B).

| Compound Name | Nav1.2 ($\mu\text{M}$) | Nav1.5 ($\mu\text{M}$) | Nav1.7 ($\mu\text{M}$) | State Dependence * |
| --- | --- | --- | --- | --- |
| Amitriptyline | 0.56 | 0.82 | 1.25 | 9.6 |
| Tetracaine | 0.26 | 0.43 | 0.28 | 15.5 |
| Vixotrigine | 1.79 | 0.93 | 1.36 | 29.8 |
| JNJ63955918 | 3.07 | 6.26 | 0.042 | 6.1 |
| PF-05089771 | 0.17 | 10.1 | 0.035 | 42.9 |
| Tetrodotoxin | 0.019 | 3.08 | 0.033 | 4.0 |
| Carbamazepine | 139.4 | 37.1 | 52.6 | 39.9 |
| Funapide | 0.51 | 0.24 | 0.46 | 2.7 |
| Mexiletine | 28.8 | 21.0 | 13.0 | 9.2 |
| Lacosamide | 392.9 | 58.5 | 104.6 | n.a. |
| Lamotrigine | 72.4 | 20.6 | 32.8 | 24.3 |
| MK-0759 | 1.50 | 1.61 | 2.05 | 18.8 |
| Lidocaine | 47.2 | 20.1 | 15.0 | 15.4 |
| TC-N 1752 | 0.14 | 1.14 | 0.046 | 10.8 |
| VX-150 | inactive | inactive | inactive | n.a |

### 3.4 Demonstration of Swarm Screening for Nav1.7 Small Molecule Inhibitors

To assess the platform readiness to support high-throughput screening, we first conducted a pilot screen using the Prestwick Chemical Library, which is a unique collection of 1520 small molecules, many of which are approved drugs (approved by the FDA, EMA, JAN, and other agencies). Before imaging, Nav1.7 spiking HEK cells were loaded with BeRST1 dye and then incubated with the compounds at 3 μM for 15 minutes. Next, the cells were stimulated with a 10 test-pulse protocol and the resulting voltage waveforms were collected for compound pharmacology evaluation. The mean spike amplitude was used to calculate the Z’ factor for each plate. All four plates displayed Z’ values higher than 0.7 and the tetracaine in-plate control wells yielded consistent IC_50_ values across all four plates, which were 0.16, 0.18, 0.24, 0.20 μM, respectively (Figure 4). The hit rate was 9.3% when 0.5 mean spike amplitude was used as the cutoff for hit selection, which is consistent with the reported frequent occurrence of Nav channel inhibition for marketed approved drugs (Zhang et al., 2014). Since many Nav channel inhibitors are not subtype-selective, we chose a set of 75 reference compounds in the Prestwick Chemical Library also with reported IC_50_ values for Nav1.5 based on automated patch clamp recordings (Harmer et al., 2011). Fifteen out of sixteen compounds with Nav1.5 IC_50_ values less than 3 μM also inhibited Nav1.7 spiking HEK signals by more than 50% indicating a lack of subtype selectivity (Supplementary Table 1). Many anti-depressant drugs are also Nav channel inhibitors (Huang et al., 2006) and the Prestwick Chemical Library identified hits which have been characterized as anti-depressant drugs, including nefazodone, doxepin, amitriptyline, nortriptyline, clomipramine, imipramine, trimipramine, and desipramine.

**Figure 4.**
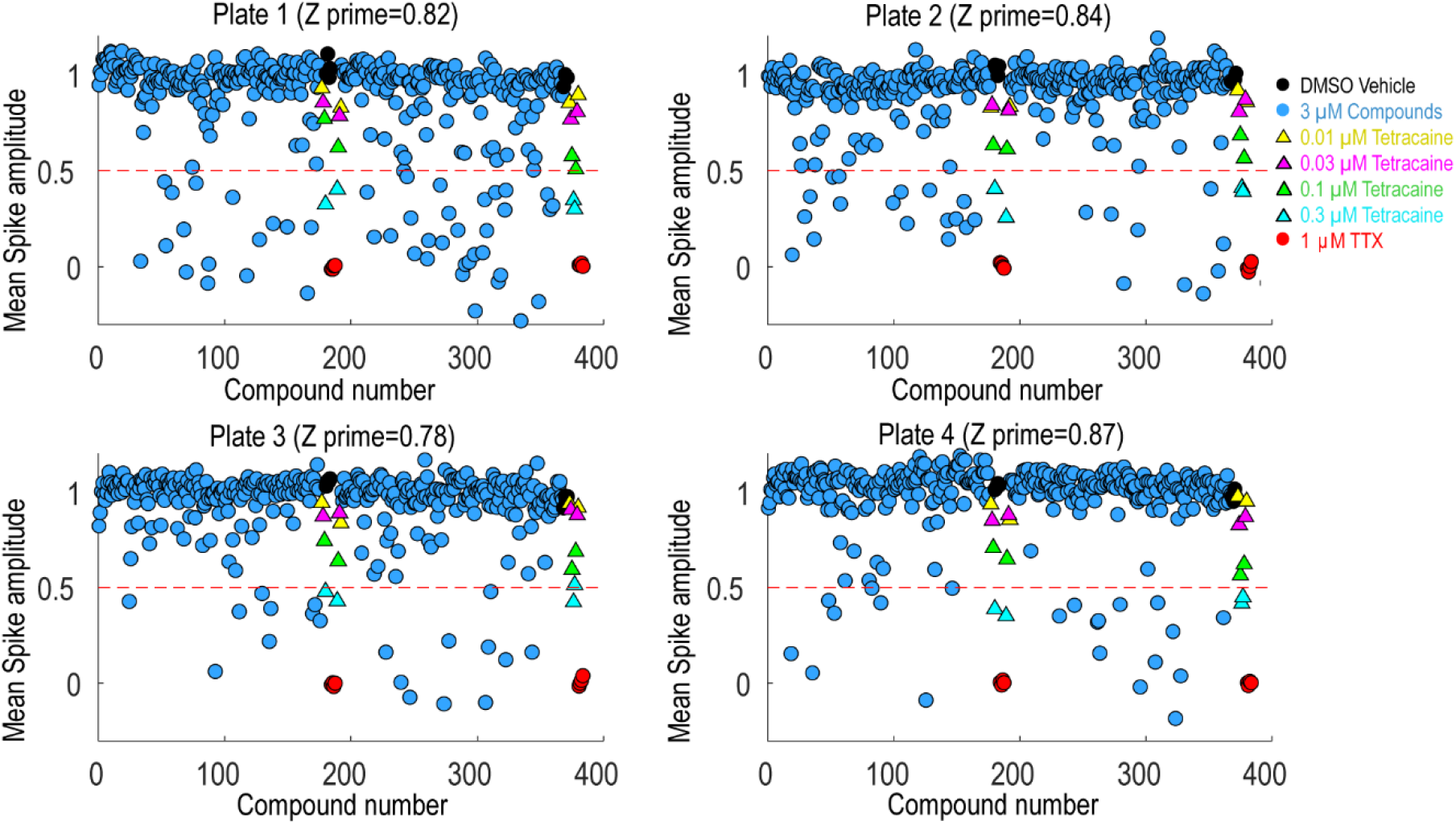
Prestwick library pilot screen. Four 384-well plates from the Prestwick library were tested using Nav1.7 spiking HEK assay under 8 mM bath potassium. The mean spike amplitude of individual wells is shown in scatter plots for each individual plate. DMSO vehicle was used as the negative control and 1 μM TTX was used as the positive control. DMSO treated wells and TTX treated wells were used to calculate the Z prime factor. On each plate, a four-dose tetracaine titration series was also included to monitor assay variability across different plates.

Following the pilot screen, we next searched for Nav1.7 inhibitors with novel chemical scaffolds and desired molecular profiles by conducting a 200,000 small molecule library screen using the Nav1.7 spiking HEK assay on the newly engineered Swarm instrument. 560 384-well plates were imaged with a throughput of ~8,000 compounds/day (Figure 5A), though 10,000 compounds/day was readily achievable. The average Z’ for the entire screen was ~0.8 (representative screening plate shown in Figure 5B) and the tetracaine in-plate control IC_50_ fluctuation was less than 4-fold (Figure 5C) demonstrating assay robustness. Using 0.45 mean spike amplitude as the cutoff for hit selection, the hit rate was ~1.4% (Figure 5D). 2,766 identified hit compounds were cherry-picked to advance into hit confirmation at 1 and 3 μM (n=2). When tested at 3 μM, the hit confirmation rate was 60% when 0.5 mean spike amplitude was applied as the cutoff (Supplementary Figure 3). Among the hits confirmed at 3 μM, 352 compounds with confirmed activity at 1 μM were selected for 8-point concentration-response curve (CRC) analysis in Nav1.7 and Nav1.5 spiking HEK assays.

**Figure 5.**
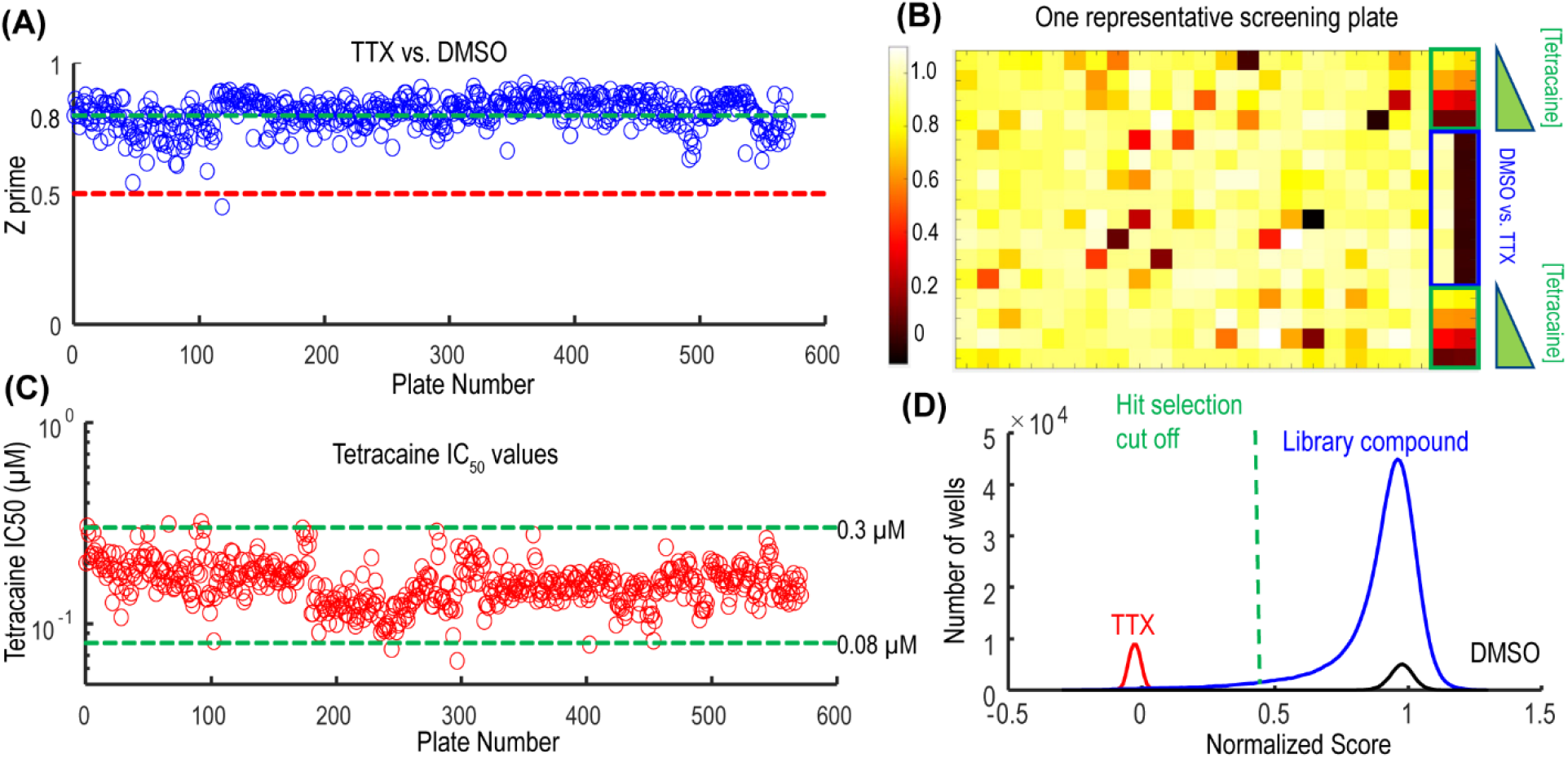
Quality control metrics analysis for 200 K Nav1.7 spiking HEK Swarm screen. (A) The Z prime factor analysis for Nav1.7 Swarm screen. All 560 plates had Z prime factors at or above 0.5. (B) The heat map of one representative screening plate. Column 1 through Column 22 contained library compounds. Columns 23 and 24 contained control compounds, including 8 wells of TTX and 8 wells of DMSO for Z prime factor calculation and two 4-point titration series of tetracaine. (C) IC_50_ values derived from the 4-point tetracaine titration for each screening plate. The IC_50_ fluctuation was less than 4-fold during the entire screen, indicating a high degree of assay consistency. (D) The histogram of library compounds and the control compounds. The normalized score of 0.45 was chosen for hit selection threshold, and based on this criterion, around 2,800 compounds were identified as hits from the 200 K Nav1.7 Swarm screen.

Validation studies were performed for 32 compounds using reordered powder samples and for activity confirmation in 10-point CRC analysis in spiking HEK assays. Reordered hit compounds were tested in Nav1.7, Nav1.5, and Nav1.2 assays for selectivity profiling and state-dependence determination (Figure 6). We benchmarked the properties of the newly identified hits against the 15 tool compounds selected in Figure 3. To mimic the genetic loss-of-function mechanism in congenital insensitivity to pain (CIP) patients, an ideal Nav1.7 inhibitor should have sufficient selectivity against both Nav1.2 and Nav1.5 and should display less strong state-dependent inhibition. Among the tested tool compounds, only JNJ63955918 demonstrated more than 10-fold subtype selectivity against both Nav1.2 (Figure 6A) and Nav1.5 (Figure 6B) and exhibited a state-dependence metric less than 10-fold (Figure 6C). However, JNJ63955918 is a small peptide that would require intrathecal delivery, which limits therapeutic applications. PF-05089771, a compound previously tested in Phase 2 clinical trials with limited success, lacked sufficient selectivity against Nav1.2 (Figure 6A) and it also displayed the strongest state-dependent block of the tested compounds (Figure 6C). Compound 1 was a promising internally identified hit from the Q-State screening library with ~10-fold subtype selectivity against Nav1.2 (Figure 6A) and more than 10-fold subtype selectivity against Nav1.5 (Figure 6B) with improved state dependent properties compared to PF-05089771. Additional structure activity relationship and exploratory medicinal chemistry efforts may lead to further improvements in subtype selectivity and reduce the level of state-dependent inhibition to achieve the desired profile for a safe and effective Nav1.7 inhibitor therapeutic for chronic pain.

**Figure 6.**
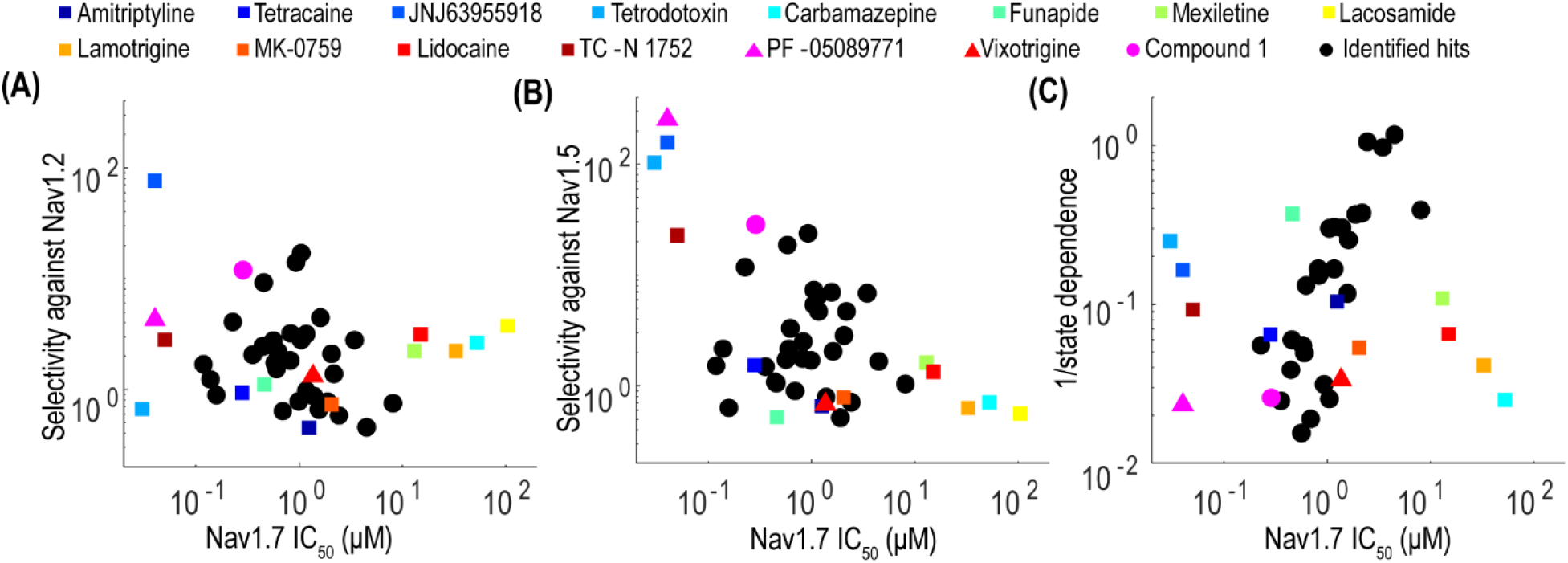
Potency, selectivity, and state dependence of tool compounds and identified hits. (A) For each Nav1.7 inhibitor, its subtype selectivity against Nav1.2, defined as the ratio of Nav1.2 IC_50_ value over Nav1.7 IC_50_ value, was plotted against its Nav1.7 IC_50_ value. (B) For each Nav1.7 inhibitor, its subtype selectivity against Nav1.5, defined as the ratio of Nav1.5 IC_50_ value over Nav1.7 IC_50_ value, was plotted against its Nav1.7 IC_50_ value. (C) For each Nav1.7 inhibitor, its reciprocal of state dependence was plotted against its potency on Nav1.7. In all three panels, the compounds with desired properties (potent, subtype selective and less state-dependence) distribute to the upper left quadrant.

### 3.5 Simultaneous Voltage and Calcium Imaging of Human iPS Cell-derived Cardiomyocytes Under Optogenetic Pacing Using the Swarm

We have previously demonstrated Optopatch compatibility with human iPS cell (hiPSC)-derived cardiomyocytes (Dempsey et al., 2016; Dempsey and Werley, 2017). Here we extended this approach by developing a Swarm-compatible counter screening assay in hiPSC-derived cardiomyocytes to assess cardiotoxicity of HTS hit compounds and validated the assay with pharmacological tool compounds. HiPSC-derived cardiomyocytes (CDI Cardiomyoctes^2^) were plated on 96-well Greiner cyclic olefin copolymer (COC) plates and transfected with blue-light activated CheRiff channelrhodopsin and calcium reporter jRGECO1a (Dana et al., 2016). Cardiomyocytes were also loaded with BeRST1 prior to imaging to record cell membrane potential. Cardiomyocytes were paced at 0.5 Hz with a ten-pulse 20 ms blue light stimulation protocol, and the BeRST1 fluorescence measure of membrane potential was recorded. For the cardiomyocyte experiments, two recordings per well were made. One recording was made prior to compound treatment and a second recording was performed following compound addition with a 10-minute compound incubation period. The post-compound treatment voltage traces were normalized to their corresponding pre-compound addition traces. Voltage traces from cardiomyocytes treated with sodium channel blockers vixotrigine, PF-05089771, and the HTS screening hit Q-State Compound 1 (QS Compound 1) are shown in Figure 7A. As a positive control, we tested the non-selective Nav channel blocker vixotrigine which showed dose-dependent reduction of the AP peak amplitude, with the AP waveform significantly distorted at the highest dose (30 μM, <25-fold above its Nav1.7 IC_50_). In comparison, the Nav channel subtype-selective PF-05089771 and QS Compound 1 caused minimal effects at the highest doses tested (3 μM and 30 μM, respectively, >40-fold above their Nav1.7 IC_50_). As additional control pharmacology, voltage traces for the calcium channel blocker nifedipine and hERG channel blockers dofetilide and E4031 were recorded (Figure 7B). Nifedipine shortened the AP width with increasing concentrations until the cardiac AP was completely extinguished at the highest 0.3 μM dose, consistent with previous reports (Dempsey et al., 2016). HERG channel blockers dofetilide and E4031 elongated APD70 with increasing concentrations, as expected (Dempsey et al., 2016; Millard et al., 2018). At the highest doses, the hERG blockers altered the AP waveforms to such a degree that the cardiomyocytes failed to fire APs at the 0.5 Hz pacing frequency. Addition of dofetilide increased the APD70 from 1s to 1.75s at 0.3 μM. Similarly, E4031 addition prolonged the AP waveform from 1s to 2.4 s at the intermediate dose of 0.03 μM.

**Figure 7.**
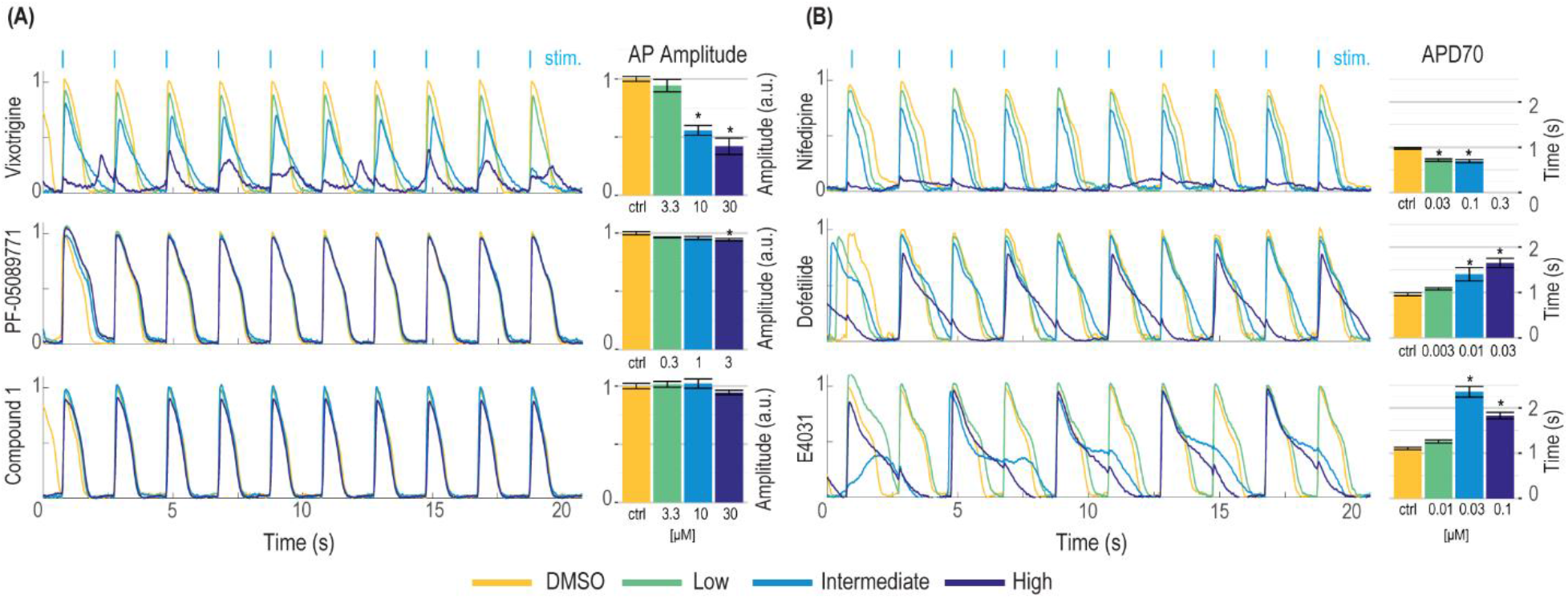
Validation of hiPSC-derived cardiomyocyte counter-screening assay on Swarm instrument with pharmacological tool compounds and an identified screening hit. Representative BeRST1 voltage fluorescence traces are shown for 0.5 Hz paced cardiomyocytes treated with increasing concentrations of compound (yellow to dark blue). Cardiomyocytes were treated with (A) sodium channel blockers (Vixotrigine, PF-05089771, QS Compound 1) and (B) calcium channel and hERG channel blockers (Nifedipine, Dofetilide, E4031). Quantification of (A) action potential (AP) peak amplitude for sodium channel blockers and (B) AP70 width for calcium channel and hERG channel blockers are shown as mean +/− SEM from triplicate wells. * = p<0.05 compared to DMSO control.

The Swarm instrument is capable of recording from up to three emission wavelengths simultaneously. This makes the instrument ideally suited for recording both the far-red BeRST1 voltage fluorescence and red jRGECO1a Ca^2+^ transients on hiPSC-derived cardiomyocytes. Figure 8A shows representative fluorescence traces overlaid with the ten-pulse blue light stimulation protocol used to optically pace the cardiomyocytes. The cardiomyocytes were treated with the calcium channel blocker nifedipine, and voltage fluorescence and calcium transients were measured simultaneously (Figure 8B). At low and intermediate doses of 0.03 μM and 0.1 μM nifedipine, respectively, the voltage peak amplitude decreased by 33%+/-2.7% and 40%+/-7.0%, respectively, until the cardiomyocytes failed to fire APs at the highest 0.3 μM dose. Similarly, the calcium transients were quantified by taking the integrated area of the calcium signal. Addition of 0.03 μM nifedipine decreased the integrated calcium area by 28% +/− 4.5% and 0.1 μM nifedipine decreased the integrated area by 31% +/− 13% until it was completely extinguished at the highest 0.3 μM dose (Figure 8C). The results show that nifedipine altered both the AP waveform and calcium transients of cardiomyocytes, as expected (Dempsey and Werley, 2017; Millard et al., 2018). The Swarm instrument was able to capture the pharmacological effects of compounds with known activities in cardiomyocytes using both voltage and calcium readouts and can provide a platform for detailed pharmacological characterization using human cardiomyocytes.

**Figure 8.**
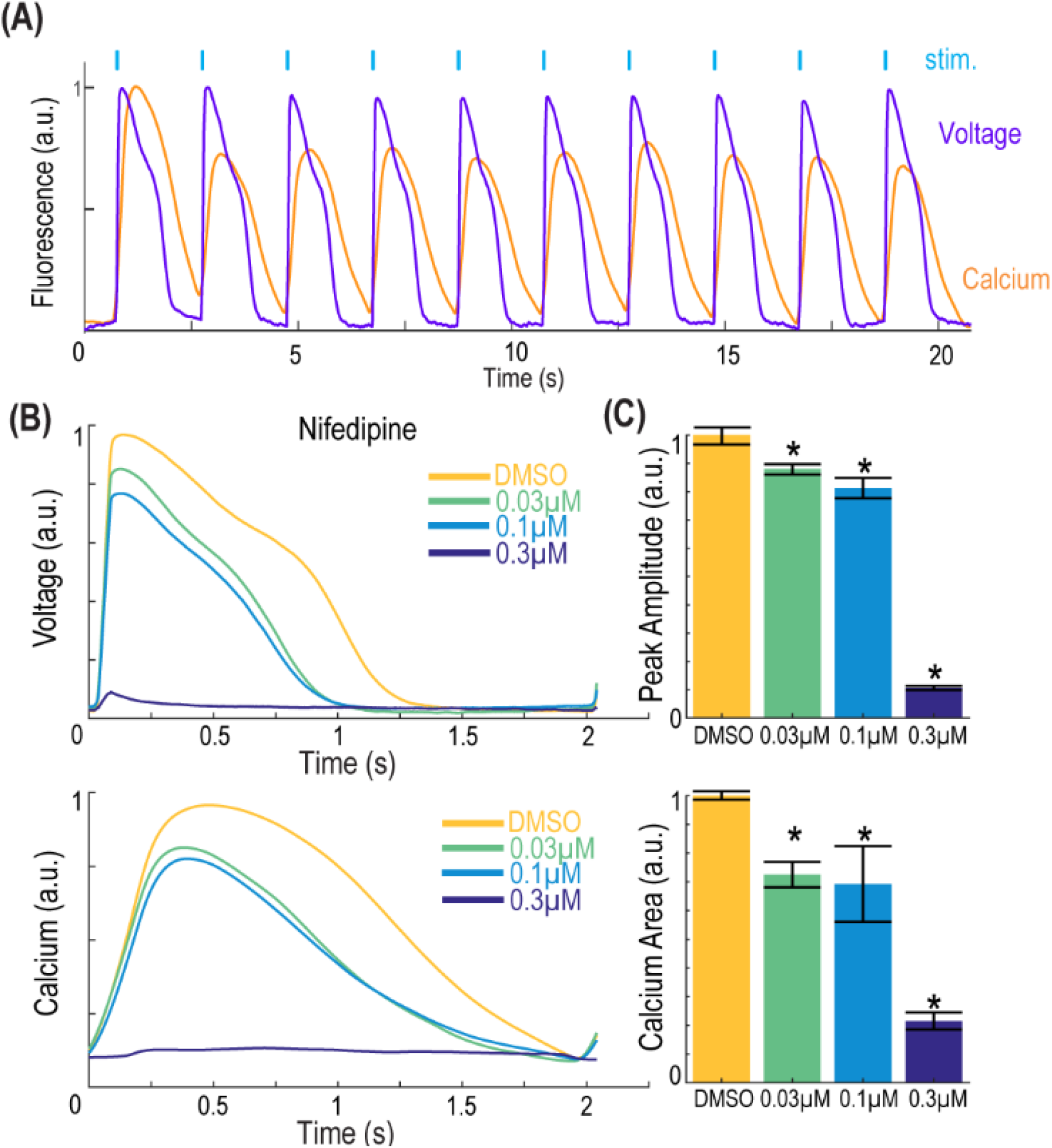
Simultaneous voltage and calcium imaging with the Swarm instrument on hiPSC-derived cardiomyocytes. (A) Representative fluorescence traces from hiPSC-derived cardiomyocytes expressing jRGECO1a reporter and loaded with BeRST1 voltage sensitive dye. Cardiomyocytes were paced with ten 0.5 Hz, 20ms blue pulses. (B) Average fluorescence waveforms for voltage and Ca^2+^ in hiPSC-derived cardiomyocytes treated with increasing concentrations of calcium channel blocker Nifedipine (yellow to dark blue). (C) Quantification of voltage AP peak amplitude and integrated Ca^2+^ waveform area. Data shown as mean from triplicate wells with +/-SEM as error bars. * = p<0.05 compared to DMSO control.

## 4 Discussion

Here we describe the design and validation of a novel optically-based instrument, the Swarm, for evaluating pharmacological effects of compounds on ion channels and excitable cells. Swarm enabled optical stimulation and simultaneous optical readout of multiple parameters including membrane voltage and intracellular calcium levels in cell-based assays. Optical stimulation via blue light stimulation of heterologously expressed channelrhodopsins allowed for precise and rapid stimulation with flexible control of stimulation protocols. Multiwavelength readouts of membrane voltage and calcium at red-shifted wavelengths enabled simultaneous stimulation and multiparameter readout with rapid response kinetics that are currently only limited by response times of the fluorescent voltage sensors. Swarm incorporated these capabilities in an instrument containing 24 individual objectives each having four optical input channels and three optical readout channels along with electronics for detection of the optical signals and control of LED-based stimulation. Swarm is designed for high throughput compound evaluation in 96- and 384-well plate formats and provides a new scalable technology for high-throughput screening of compounds targeting ion channels and excitable cells.

Optical assays provide an advantageous path to develop scalable high throughput assays. Specialized and expensive consumable items are not typically required, and the screening format can be readily changed to match assay specific needs and throughput. Previous assays methods used for sodium channel screening have relied on flux of permeant ions (Trivedi et al., 2008; Du et al., 2015), fluorescence measurements of membrane potential (Dyes et al., 2004), and automated electrophysiology (Chambers et al., 2016; Li et al., 2017; Zhang et al., 2020). Flux assays and fluorescent membrane potential assays can provide a robust measure of cumulative sodium channel activity and can be tuned to detect specific pharmacological profiles, but they lack temporal resolution and generally require use of non-physiological triggering agents, such as veratridine, which can alter the pharmacological sensitivity of the assay. Binding assays typically use radiolabeled ligands which are less amenable to HTS and the readouts are not directly coupled to sodium channel function. Automated electrophysiology instruments provide a high-resolution linear readout of sodium channel function, but consumable costs and available assay formats can limit use in HTS campaigns and position this technology as best used for medium throughput assays and downstream secondary confirmation of compound activity.

The optogenetic assays implemented on Swarm provide the throughput and scalability of conventional fluorescent sodium channel assays such as FLIPR coupled with the information content, temporal resolution, activation by a physiological stimulus, and flexible control of channel activation obtained with automated E-Phys. In this work, we demonstrate the capabilities of Swarm to execute a Nav1.7 spiking HEK HTS campaign and to perform selectivity assays using Nav1.2 and Nav1.5 spiking HEK assays and hiPSC-derived cardiomyocyte counter screens along with a mechanistic classification of hit compounds. The subtype selectivity, state dependence, and cardiomyocyte toxicity information obtained from Swarm assays provide pharmacological profiles that can be readily generated and contain key information to drive chemical optimization efforts and to prioritize compound selection. The profiles generated for compounds listed in Figure 3 and shown in Figure 6 provide potential explanations for why some clinical stage compounds (e,g, vixotrigine and PF-05089771) demonstrated limited clinical success due to a lack of subtype selectivity (vixotrigine) or highly state-dependent inhibition (PF-05089771). The Swarm instrument provided a stable and reliable platform to execute a large-scale screen and downstream activities. We have also demonstrated that spiking HEK assays on Swarm provided reliable compound potency estimates of well-known reference compounds that matched values from previously published literature and from a similar assay performed using a single objective single well serial epifluorescence microscope (Zhang et al., 2020). Future developments of the instrument will implement laser-based illumination to enable the required intensities to fully leverage optogenetic voltage reporters coupled with novel sensing devices to enable extension to 1536-well plate formats for yet higher throughput screening along with new multiplexing capabilities.

Drug discovery projects require integrated use of a wide array of biophysical, biochemical, cell-based, and in vivo measurement modalities. Advances in drug discovery are often enabled by technological advances, and technology development can be motivated by drug discovery progress in a virtuous cycle. This relationship was highlighted more generally by Sidney Brenner, “Progress in science depends on new techniques, new discoveries and new ideas, probably in that order” (Brenner, 2002). More specifically, drug discovery for ion channels and excitable cells such as neurons and cardiomyocytes has been propelled by technical developments including patch clamp recordings (Hamill et al., 1981), fluorescence plate readers (Schroeder, 1996; Huang et al., 2006), and automated patch clamp (Fertig et al., 2002; Schroeder et al., 2003). In general, choice of assay methods aims to optimize information content, throughput, and relevance to disease pharmacology (McManus, 2014). Optogenetic assays implemented using the Swarm instrument enable a novel system that combines high information content, high throughput, and deep pharmacological characterization which can be leveraged to drive diverse therapeutic discovery programs for nervous system disorders and other disease areas involving excitable cells.

## 5 Conflict of Interest

GBB, HZ, BNH, JJ, JG, RC, DZ, AB, CW, YL, SN, OBM and GTD are current or former employees of Q-State Biosciences and may hold stock options in Q-State Biosciences.

## 6 Author Contributions

GBB, HZ, BNH, CW, SN, OBM and GTD designed the study. GBB, YL, AB and SN built the instrument and acquisition software. GBB, HZ, BNH, JJ, JG, RC and DZ acquired the data. GBB, HZ, and BNH analyzed the data. GBB, HZ, BNH, YL, OBM and GTD interpreted the results. GBB, HZ, BNH, YL, OBM and GTD prepared the manuscript. All authors contributed to manuscript revision and approved the submitted version.

## 7 Funding

This work was supported by funding from National Heart, Lung and Blood Institute grant number 2R44HL126314-02A1 and National Institute of Neurological Disorders and Stroke grant number 1R44NS107041-01.

## 6 Acknowledgements

We would like to thank Steve Wasserman for input on instrumentation engineering and David Gerber for input on the manuscript.

## 10 Permission to Reuse and Copyright

Arrow-wavy-right-medium, sodium-potassium-pump-membrane-ions, and 7helix-receptor-membrane by Servier https://smart.servier.com/ are licensed under CC-BY 3.0 Unported https://creativecommons.org/licenses/by/3.0/ and were adapted for Figure 2A.

