## Supplementary Information for "Highly Parallelized, Multicolor Optogenetic Recordings of Cellular Activity for Therapeutic Discovery Applications in Ion Channels and Disease-associated Excitable Cells"

### *Supplementary Material*

#### 1 Supplementary Figures and Tables

##### 1.1 Supplementary Figures

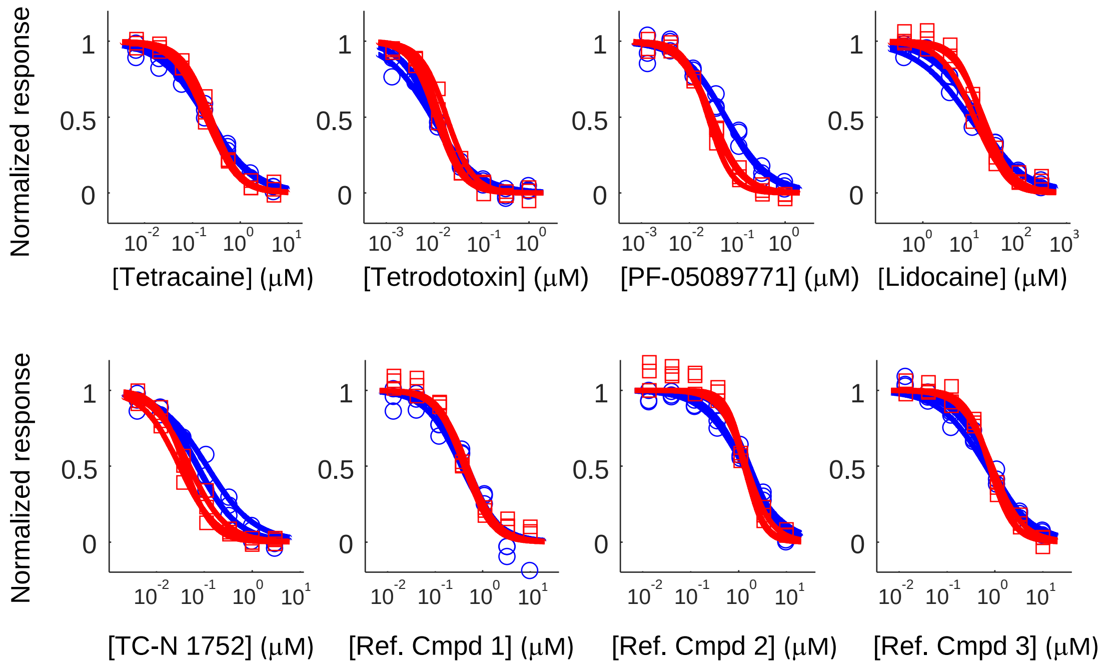


**Supplementary Figure 1. Cross comparison between Swarm and epifluorescence microscope spiking HEK assay.** Concentration response curves of 5 Nav channel tool compounds (Amitriptyline, Tetracaine, Tetrodotoxin, PF-05089771, Lidocaine, TC-N 1752) and 3 internal identified reference compounds determined using. Nav1.7 spiking HEK assay imaged with the Swarm instrument (blue curves) or with a custom-built, single well epifluorescence Optopatch microscope (red curves). Each compound has three titration series from the same compound plate.

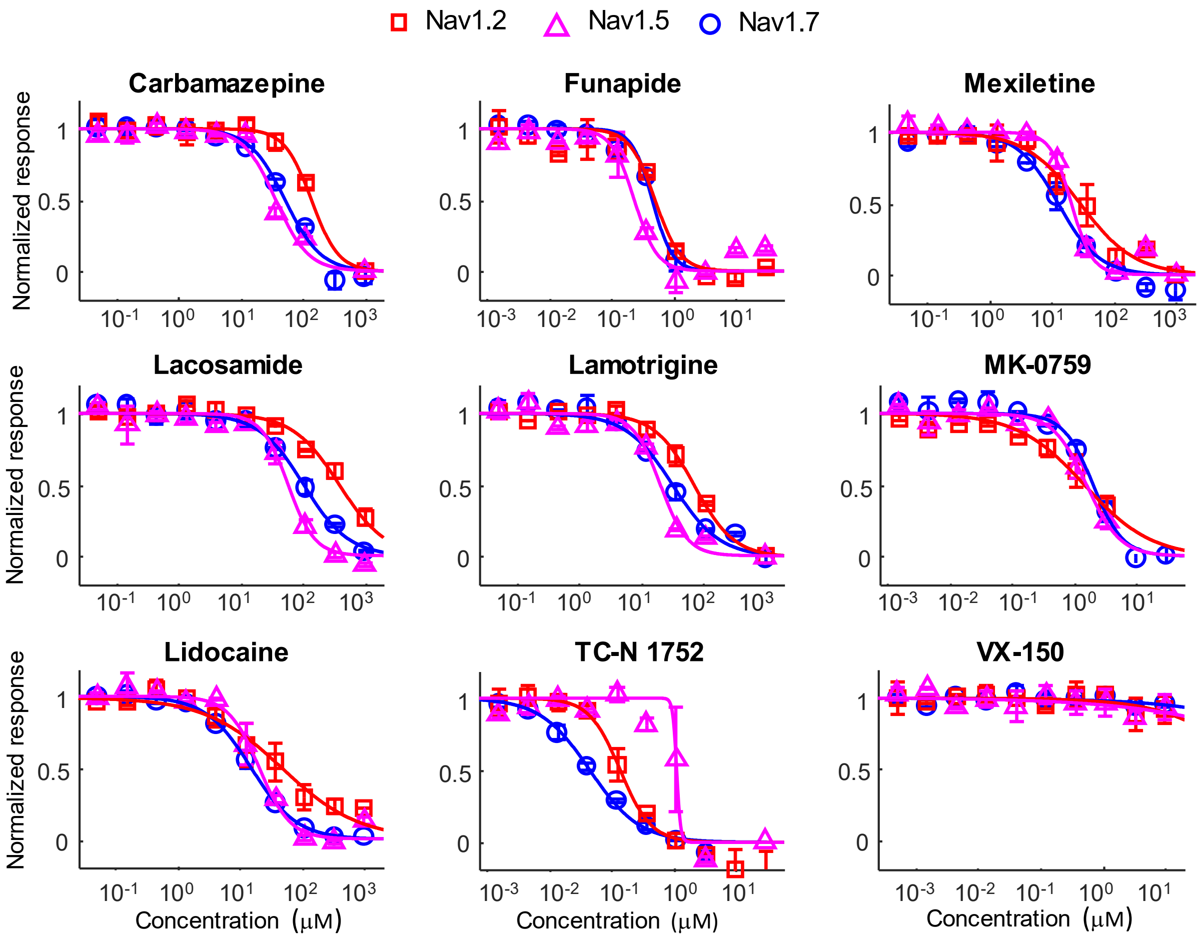


**Supplementary Figure 2. Tool compound subtype selectivity profiles.** Subtype selectivity of 9 additional Nav channel tool compounds. In addition to the 6 compounds shown in Figure 3, 9 additional tool compounds were tested on Nav1.2 (8 mM bath [$K^+$]), Nav1.5 (6 mM bath [$K^+$]), and Nav1.7 (8 mM bath [$K^+$]) using Swarm spiking HEK assays. The $IC_{50}$ value fitting results can be found in Table 1.

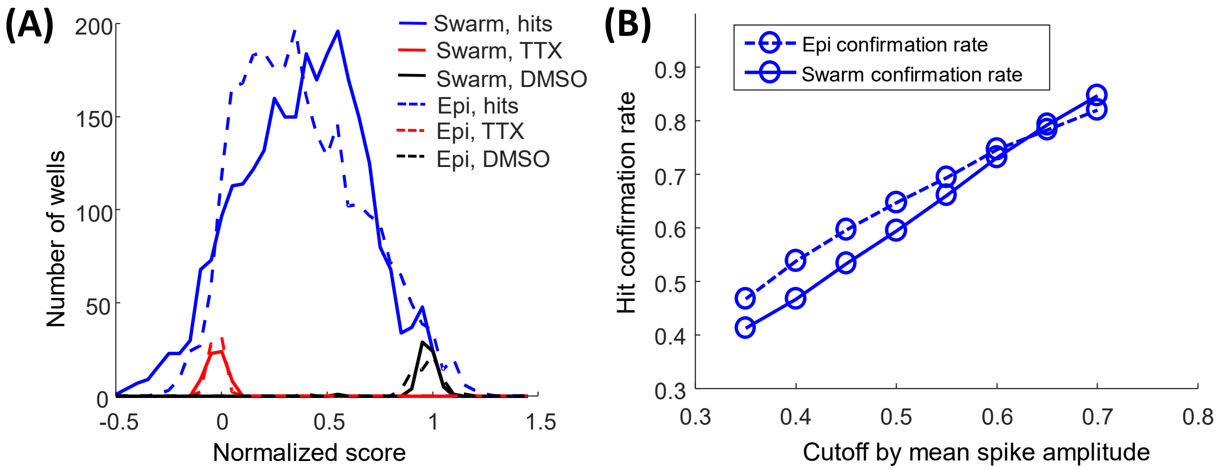


**Supplementary Figure 3**. **Swarm spiking HEK 200K compound screen hit confirmation results at 3 μM.** Around 2,800 hits identified from a 200K compound Swarm screen were cherry-picked from the library and retested at 3 μM in duplicate. (A) The cherry-picked hits were tested in both the Swarm instrument and the epifluorescence microscope. The distribution of normalized scores based on mean spike amplitude from both the Swarm instrument and the epifluorescence microscope are shown in (A). TTX at 1 μM was included as the positive control and DMSO was included as the negative control. (B) The hit confirmation rate of the cherry-picked hits from either Swarm instrument or the epifluorescence microscope was plotted against the exact cutoff used to define the confirmed hits.

#### 1.2 Supplementary Tables

| Compound | Nav1.7 Swarm (score at 3 μM) | Nav1.5 Ionworks ($IC_{50}$, μM) | Compound | Nav1.7 Swarm (score at 3 μM) | Nav1.5 Ionworks ($IC_{50}$, μM) |
|---|---|---|---|---|---|
| Moricizine | 0.15 | 1.2 | Quinidine | 0.76 | 9.7 |
| Propafenone | 0.20 | 1.2 | Amodiaquin | 0.95 | 9.7 |
| Amitriptyline | 0.16 | 1.6 | Reserpine | 1.01 | 10.5 |
| Pimozide | 0.89 | 1.7 | Clozapine | 0.74 | 11.6 |
| Bepridil | -0.28 | 1.9 | Diltiazem | 0.82 | 14.2 |
| Nortriptyline | 0.19 | 2.0 | Citalopram | 0.92 | 14.7 |
| Terfenadine | 0.39 | 2.0 | Ambroxol | 0.80 | 15.2 |
| Haloperidol | -0.09 | 2.3 | Diphenhydramine | 0.88 | 16.4 |
| Astemizole | 0.01 | 2.3 | Riluzole | 0.14 | 17.6 |
| Nilvadipine | 0.24 | 2.3 | Dipyridamole | 0.92 | 19.3 |
| Desipramine | 0.38 | 2.4 | Risperidone | 0.89 | 20.2 |
| Flunarizine | 0.16 | 2.6 | Quinacrine | 0.96 | 20.2 |
| Clomipramine | 0.20 | 2.6 | Pyrimethamine | 0.99 | 24.6 |
| Trimipramine | 0.37 | 2.7 | Mexiletine | 1.00 | 25.7 |
| Chlorpromazine | 0.32 | 2.8 | Mifepristone | 0.97 | 25.8 |
| Loperamide | 0.04 | 2.9 | Lofexidine | 0.88 | 27.2 |
| Protriptyline | 0.63 | 3.1 | Lidocaine | 0.91 | 30.9 |
| Perhexiline | 0.59 | 3.4 | Hydroxyzine | 0.56 | 32.8 |
| Thioridazine | 0.04 | 3.5 | Nifedipine | 0.97 | 33.5 |
| Imipramine | 0.35 | 3.6 | Loratadine | 0.62 | 33.6 |
| Clemastine | 0.02 | 4.0 | Chloroquine | 0.98 | 36.5 |
| Nicardipine | 0.07 | 4.3 | Olanzapine | 1.09 | 39.0 |
| Bupivacaine | 0.27 | 4.3 | Bromopride | 0.92 | 41.4 |
| Sertraline | 0.87 | 4.3 | Ketoconazole | 0.96 | 44.9 |
| Amiodarone | 0.52 | 4.8 | Lamotrigine | 0.99 | 63.4 |
| Fluoxetine | 0.64 | 4.9 | Exemestane | 0.90 | 71.3 |
| Domperidone | 0.53 | 5.6 | Prilocaine | 0.95 | 72.6 |
| Flecainide | 0.72 | 5.8 | Bupropion | 0.97 | 76.2 |
| Amoxapine | 0.88 | 5.8 | Mepivacaine | 0.96 | 81.4 |
| Cyproheptadine | 0.44 | 6.1 | Procaine | 1.07 | 84.6 |
| Salmeterol | 0.22 | 7.2 | Venlafaxine | 0.95 | 90.1 |
| Mesoridazine | 0.40 | 7.2 | Memantine | 0.98 | 92.6 |
| Trifluoperazine | 0.25 | 7.4 | Buspirone | 0.97 | 125.4 |
| (R)-Propranolol | 0.63 | 7.5 | Ziprasidone | 1.07 | 170.0 |
| Fluphenazine | 0.31 | 8.0 | Moxifloxacin | 1.06 | 206.7 |
| Sertindole | 0.64 | 8.1 | Chloramphenicol | 1.07 | 215.0 |
| Verapamil | 0.38 | 9.3 | Disopyramide | 0.94 | 302.3 |
| | | | Procainamide | 1.00 | 2050.0 |

**Supplementary Table 1. Potencies of Prestwick library compounds in Nav1.7 Swarm assay and reported Nav1.5 automated patch clamp assay.** 75 compounds with literature reported Nav1.5 IC50 values were ranked based on their potencies. Their potencies in Nav1.7 Swarm spiking HEK assay were provided in the same table. The compounds that were identified as hits in the Prestwick library screen are highlighted in green.
